# Human-associated microbiota suppress invading bacteria even under disruption by antibiotics

**DOI:** 10.1101/2020.09.02.279307

**Authors:** Andrew D. Letten, Michael Baumgartner, Katia R. Pfrunder-Cardozo, Jonathan Levine, Alex R. Hall

**Affiliations:** School of Biological Sciences, University of Queensland, Brisbane, Queensland 4072, Australia; Institute of Integrative Biology, Department of Environmental Systems Science, ETH Zürich, 8092 Zürich, Switzerland; Dept of Ecology and Evolutionary Biology, Princeton University, Princeton, NJ, 08544-1003 USA

## Abstract

In light of their adverse impacts on resident microbial communities, it is widely predicted that broad-spectrum antibiotics can promote the spread of resistance by releasing resistant strains from competition with other strains and species. We investigated the invasion success of a resistant strain of *Escherichia coli* inoculated into human-associated communities in the presence and absence of the broad and narrow spectrum antibiotics rifampicin and polymyxin B, respectively. We found strong evidence of community-level suppression of the resistant strain in the absence of antibiotics and, despite large changes in community composition and abundance following rifampicin exposure, suppression of the invading resistant strain was maintained in both antibiotic treatments. Instead, the strength of competitive suppression was more strongly associated with the individual donor from which the community was sampled. This suggests microbiome composition strongly influences susceptibility to invasion by antibiotic-resistant strains, but at least some antibiotic-associated disruption can be tolerated before invasion susceptibility increases. A deeper understanding of this association will aid the development of ecologically-aware strategies for managing antibiotic resistance.

---

The overuse of broad-spectrum antibiotics in clinical and agricultural settings is a key driver of the current antibiotic resistance crisis [1]. Research into antibiotic resistance has traditionally focused on the evolution of resistance in individual pathogens [2]. In the last decade, researchers have turned their attention to the collateral damage inflicted on commensal members of the microbiome, such as those belonging to the dense communities of the human gastrointestinal tract [3, 4]. Several studies have shown that antibiotics can leave gut communities vulnerable to colonisation by other pathogens [5–7], and, most recently, resistance evolution in invading strains can be facilitated by the absence of community suppression [8, 9]. Taken together, these two lines of enquiry appear to bear out conventional wisdom that relative to narrow-spectrum antibiotics or antibiotic-free conditions, broad spectrum antibiotics should increase the likelihood of communities being invaded by resistant strains [10, 11]. On the other hand, given evidence that community-level properties can sometimes be robust to changes in taxonomic composition [12], it is possible that some antibiotic-associated disruption can be tolerated before colonization resistance is affected. Despite the importance of these contrasting predictions, there have been few, if any, direct tests in human-associated micro-biota.

We investigated the effect of broad and narrow spectrum antibiotics on the strength of competitive suppression on a resistant variant of a focal strain (*Escherichia coli* K-12 MG1655) inoculated into gut microbiome communities collected from human faecal samples. The focal strain was jointly resistant to the broad-spectrum antibiotic rifampicin (targets gram-positives and gram-negatives) and the narrow spectrum antibiotic polymyxin B (only targets gram-negatives). The focal strain was inoculated alongside live or sterile slurry obtained from one of three healthy human donors (described in [9]) into customized gut media without antibiotics or supplemented with 128 *µ*g/ml rifampicin or 4 *µ*g/ml polymyxin B. Following 24 hours growth under anaerobic conditions, focal strain density and total biomass were measured via colony counting and flow cytometry, and community composition and diversity were analysed via 16S rRNA sequencing.

In the absence of either antibiotic, focal strain density after 24 hours was significantly lower in the presence of the three donor communities, indicative of strong competitive suppression (Fig. 1a). Surprisingly, we detected similarly strong competitive suppression in both the antibiotic treatments as we did in the antibiotic-free treatment. Instead, we found that focal strain performance was a stronger function of the specific donor community, irrespective of antibiotic treatment (Fig. 1b).

**Figure 1:**
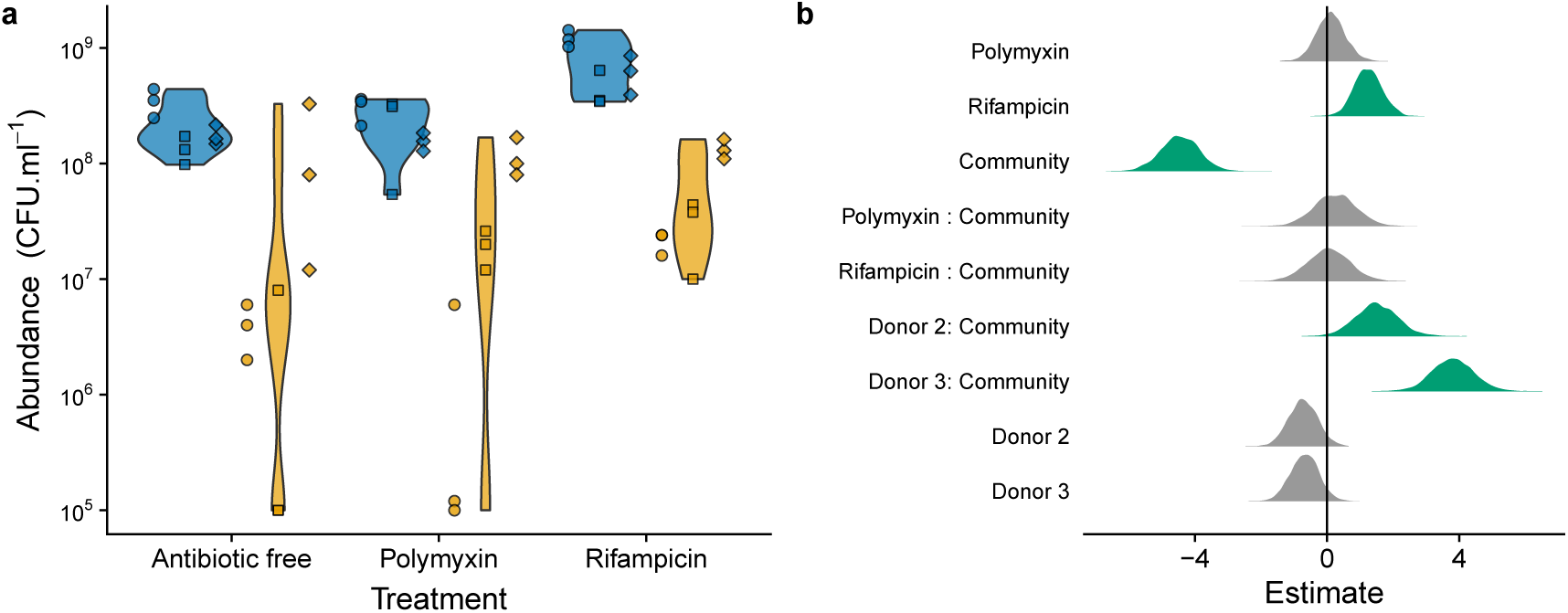
(**a**) Abundance of focal strain in each antibiotic treatment. Blue denotes community free treatments; yellow denotes community treatment. Point shape denotes community/slurry donor: donor 1 = circles, donor 2 = squares, donor 3 = diamonds. **(b)** Model coefficients (posterior distributions) from a linear model (negative binomial errors) of focal strain abundance as a function of community, antibiotic, and donor, and the interactions between community and antibiotic, and community and donor. Posteriors in green have 95% credible intervals that do not overlap with 0 (i.e., there is less than 5% probability there is no effect of the variables/interactions captured by these coefficients). Intercept (not shown) = Donor 1 in the no antibiotic treatment in the absence of the community (i.e. sterilized slurry)

What makes these results particularly striking is that, consistent with previous studies [7, 10, 13], treatment with a broad-spectrum antibiotic was still associated with a marked shift in community composition (analysis of 16S amplicon data) (Fig. 2a). Based on OTU composition, all three donors in the rifampicin treatment cluster separately from the polymyxin B and antibiotic-free treatments, which cluster together (Fig. 2b). This divergence in composition appears to be largely driven by enrichment of both Enterobacteriaceae and Erysipelotrichaceae in the rifampicin treatment (Fig. 2a). In addition to strong shifts in composition, total bacterial abundance was significantly reduced in the rifampicin treatment (Fig. 2c and Fig. S2). Despite this, total richness and Shannon diversity after 24 hours did not differ between the treatments (Fig. 2c). In contrast, diversity loss over time was more strongly associated with donor identity, with the donor community associated with the weakest competitive suppression (donor 3) also exhibiting the largest decline in richness and diversity across all treatments.

**Figure 2:**
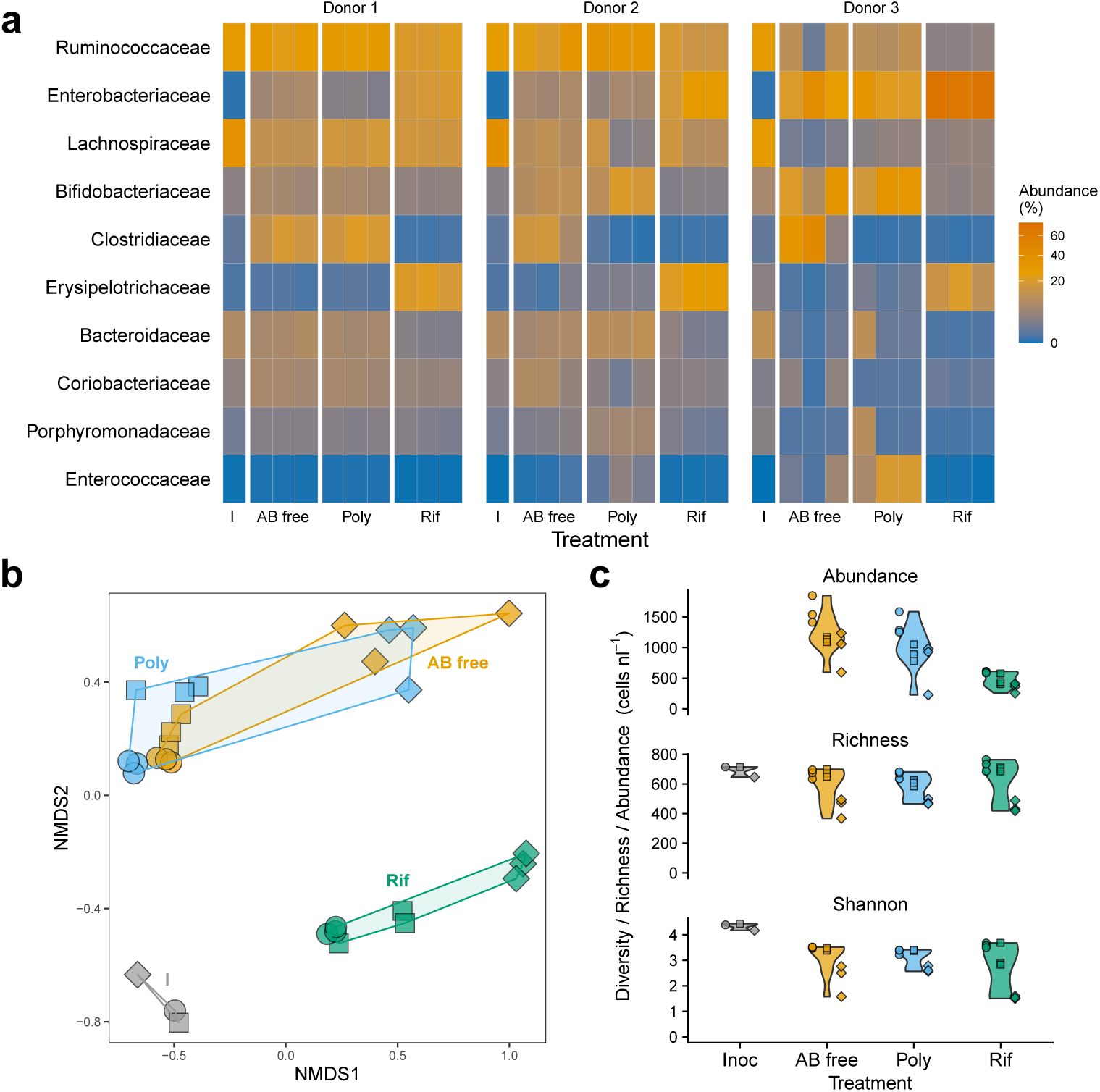
(**a**) Heatmap of relative abundance of the ten most abundant families of bacteria across treatments (derived from amplicon data). I = inoculum; AB free = Antibiotic free; Poly = polymyxin B; Rif = rifampicin. (**b**) NMDS ordination of family level composition in each treatment-donor combination. (**c**) Abundance (top), species richness (middle) and Shannon diversity (bottom) in each treatment. In **b** & **c**: circles = donor 1; squares = donor 2, diamonds = donor 3.

A limitation of this study is that we only considered the effects of two antibiotics. Nevertheless, given the scale of community perturbation observed (Fig. 2), we can at least be sure our findings do not stem from a weak treatment effect. There must be some limit dictated by antibiotic concentration, combination, or duration of exposure, beyond which we would expect to observe stronger competitive release. Indeed, prior research has shown that antibiotics can greatly inhibit colonisation resistance [14, 15]. As such, characterizing where this limit lies will be an important challenge for future work. Similarly, although we only considered a single focal strain, and other strains/species may have been more or less invasive, key for our experiment was that the focal strain had a positive growth rate over the timescale of the experiment, despite exhibiting significant resistance costs in antibiotic-free assays (Fig. S1). This allowed us to test for sensitivity of invasion success to antibiotic treatment. We also note that in spite of a small unanticipated boost in the focal strain’s performance in the presence of rifampicin in the absence of the community (a possible hormetic response [16] absent under aerobic growth in LB, Fig S1), we did not observe an increase in the magnitude of competitive release in the rifampicin treatment. Finally, the drop in Shannon diversity indicates, unsurprisingly, microcosms are a novel environment relative to the source environment. Despite this, key taxa in each community were stable over the course of the experiment, and previously over a longer timescale in the same setup [9], demonstrating these conditions sustain diverse human-associated communities over relevant timescales.

In conclusion, on the one hand, these results are entirely consistent with prevailing wisdom that healthy gut communities can suppress invading strains and thereby reduce the likelihood of resistance emerging [8, 9, 17]. On the other hand, the absence of a significant effect of broad, or even narrow, spectrum antibiotics on the degree of competitive suppression of our focal strain is much more surprising. This shows that the functional diversity of gut communities may be more robust to disturbance by broad spectrum antibiotics than previously recognised. This is not to suggest that the use of broad-spectrum antibiotics does not drive marked changes in composition but rather that there is some degree of functional redundancy in diverse communities that facilitates the maintenance of competitive suppression [12, 18]. Notwithstanding the need to test how these findings translate to *in vivo* settings, this finding is relevant for optimizing personalised treatments that either account for disruption by antibiotics or that make microbiomes harder for pathogens to invade.

## Acknowledgements

This project received funding from the European Union’s Horizon 2020 research and innovation programme under the Marie Skłodowska-Curie grant agreement No 750779. AH acknowledges Swiss National Science Foundation Project 31003A 165803. We thank the Genetic Diversity Centre (GDC) ETH Zurich for sequencing and bioinformatics support.

## Supporting Information

Additional supporting information may be found in the online version of this article:

## References

[1] Blair, J. M., Webber, M. A., Baylay, A. J., Ogbolu, D. O. & Piddock, L. J. Molecular mechanisms of antibiotic resistance. Nature Reviews Microbiology 13, 42–51 (2015).

[2] Palmer, A. C. & Kishony, R. Understanding, predicting and manipulating the genotypic evolution of antibiotic resistance. Nature Reviews Genetics 14, 243–248 (2013).

[3] Blaser, M. J. Antibiotic use and its consequences for the normal microbiome. Science 352, 544–545 (2016).

[4] Modi, S. R., Collins, J. J. & Relman, D. A. Antibiotics and the gut microbiota. Journal of Clinical Investigation 124, 4212–4218 (2014).

[5] Lawley, T. D. & Walker, A. W. Intestinal colonization resistance. Immunology 138, 1–11 (2013).

[6] Libertucci, J. & Young, V. B. The role of the microbiota in infectious diseases. Nature Microbiology 4, 35–45 (2019).

[7] Bhalodi, A. A., van Engelen, T. S. R., Virk, H. S. & Wiersinga, W. J. Impact of antimicrobial therapy on the gut microbiome. Journal of Antimicrobial Chemotherapy 74, i6–i15 (2019).

[8] Klümper, U. et al. Selection for antimicrobial resistance is reduced when embedded in a natural microbial community. The ISME Journal 1–11 (2019).

[9] Baumgartner, M., Bayer, F., Pfrunder-Cardozo, K. R., Buckling, A. & Hall, A. R. Resident microbial communities inhibit growth and antibiotic-resistance evolution of Escherichia coli in human gut microbiome samples. PLOS Biology 18, e3000465 (2020).

[10] Ianiro, G., Tilg, H. & Gasbarrini, A. Antibiotics as deep modulators of gut microbiota: Between good and evil. Gut 65, 1906–1915 (2016).

[11] Spaulding, C. N., Klein, R. D., Schreiber, H. L., Janetka, J. W. & Hultgren, S. J. Precision antimicrobial therapeutics: The path of least resistance? npj Biofilms and Microbiomes 4, 1–7 (2018).

[12] Moya, A. & Ferrer, M. Functional Redundancy-Induced Stability of Gut Microbiota Subjected to Disturbance. Trends in Microbiology 24, 402–413 (2016).

[13] Jernberg, C., Löfmark, S., Edlund, C. & Jansson, J. K. Long-term impacts of antibiotic exposure on the human intestinal microbiota. Microbiology 156, 3216–3223 (2010).

[14] Van Der Waaij, D., Berghuis-de Vries, J. M. & Lekkerkerk-Van Der Wees, J. E. Colonization resistance of the digestive tract in conventional and antibiotic-treated mice. Journal of Hygiene 69, 405–411 (1971).

[15] Kim, S., Covington, A. & Pamer, E. G. The intestinal microbiota: Antibiotics, colonization resistance, and enteric pathogens. Immunological Reviews 279, 90–105 (2017).

[16] Mathieu, A. et al. Discovery and Function of a General Core Hormetic Stress Response in E. coli Induced by Sublethal Concentrations of Antibiotics. Cell Reports 17, 46–57 (2016).

[17] Pamer, E. G. Resurrecting the intestinal microbiota to combat antibiotic-resistant pathogens. Science 352, 535–538 (2016).

[18] Louca, S. et al. Function and functional redundancy in microbial systems. Nature Ecology and Evolution 2, 936–943 (2018).

