## Supplementary Information for "Human-associated microbiota suppress invading bacteria even under disruption by antibiotics"

September 2, 2020

This file includes:

Material and methods
Supplementary Figure S1-S2

*These authors contributed equally.

### Materials and Methods

#### Focal strain

We used an *Escherichia coli* K-12 MG1655 strain with a yfp::ampicillin resitance cassette and two point mutations conferring resistance against polymyxin (*basS*) and rifampicin (*rpoB*) as our focal strain. One day prior to the microcosm experiment, we inoculated *E.coli* in LB and incubated it overnight at 37°C in a shaking incubator.

#### Microcosm experiment

We used stool samples from consenting human donors that we collected on the 15th of May 2018. The sampling protocol, approved by the ETH Zürich Ethics Commission (EK 2015-N-55) is described in [1]. In brief, we collected samples from three anonymous, healthy human donors and mixed the samples with 200 ml of anaerobic peptone wash (1 g/l peptone, 0.5 g/l L-Cysteine, 0.5 g/l bile salts and 0.001 g/l Resazurin; Sigma-Aldrich) to make a 10% (w/v) anaerobic faecal slurry. We divided the faecal slurry in two equal volumes and autoclaved one of the aliquots to create sterilized (community-free) slurry and stored it at -80°C. We added to the other half of the aliquot glycerol as a cryo-protective agent and stored it at -80°C until the day of the experiment. There is a risk that freeze-thawing could lead to the loss of some taxa from the sampled microbiota [2], but previous studies indicate this risk is small, with frozen slurry containing rich, viable communities [3–5].Note also that we do not exclude loss of some taxa, but for this experiment the aim was to capture abundant and rich communities sampled from humans (not to recover 100% of all species present in the GI tract).

Prior to the microcosm experiment, we filled 54 Hungate tubes with 7.2 ml basal medium (2 g/l Peptone, 2 g/l Tryptone, 2 g/l Yeast extract, 5 g/l NaCl, 0.04 g K2HPO4, 0.04 g/l $KH_2PO_4$, 0.01 g/l $MgSO_4x7H_2O$, 0.01 g/l $CaCl_2x6H_2O$, 0.005 g/l Hemin, 0.5 g/l L-Cysteine, 2g/l Starch, 1.5 g/l casein, 0.001g/l Resazurin, pH adjusted to 7; Sigma-Aldrich) flushed the headspace with nitrogen to remove oxygen, sealed and autoclaved the tubes. We thawed sterile and live slurries from each human donor on the morning before the experiment. For each human donor background, we inoculated $2x10^6$ of focal strain with the resident microbial community (n=9), containing 350 $\mu$l of live faecal plus 500 $\mu$l of the sterilized slurry, and in absence of the community, containing 850 $\mu$l of sterilized slurry (community-free treatment, n=9). To each of these tubes we added either rifampicin (128 $\mu$g/ml), polymyxin (4 $\mu$g/ml) or no antibiotics as a control treatment. Before and after incubation at 37°C in a shaking incubator for 24h, we took samples for plating, flow cytometry and sequencing. For plating, we diluted samples in PBS (Phosphate Buffered Solution) and plated them on chromatic MH agar (Liofilchem) supplemented with rifampicin to count abundance of our focal strain.

We quantified total bacterial densities by flow cytometry (Novocyte 2000R, ACEA Biosciences Inc.). We diluted each sample containing live community with PBS, stained them with Sybrgreen (final concentration of $10^{-4}$ of the stock solution, Life Technologies, Zug, Switzerland) and detected bacteria based on their signature in a cytogram of green fluorescence versus forward scattered light. For sequencing, we took samples from all microbiome treatments at 0 and 24h, supplemented them with glycerol and stored them at -80°C until further processing.

#### Amplicon sequencing

We used the powerlyzer power soil kit (Qiagen) to extract DNA for amplicon sequencing according to the manufacturers protocol with the following modification. We concentrated each sample before extraction by adding 1.5 ml of each sample in an extraction tube provided by the kit, centrifuged for 10 min at 10000 rpm, discarded the supernatant and repeated this step. DNA quality and quantity was assessed by Nanodrop and Qubit.

We sequenced the variable regions V3-V4 of the 16S rRNA gene with previously described primers [6]. Following the Illumina 16S Metagenomic Sequencing Library preparation guide for the MiSeq Illumina sequencing platform we amplified target regions with a limited 17 cycle PCR, followed by a PCR clean-up and adapter attachment of the Nextera XT Index Kit v2 in a second PCR step. We assessed quality and quantity of amplicon library by TapeStation and qPCR using the KAPA library quantification kit. After normalization and pooling we sequenced the library on the MiSeq sequencing platform using 2x300 V3 kit. We used Trimmomatic and FLASH to demultiplex and quality-filter the raw paired-end sequencing reads. We used Usearch software to cluster ZOTUs with a 99% similarity threshold. For taxonomic assignment prediction against the Silva 16S rRNA database we used SINTAX.

#### Statistical analysis

Generalised linear models for both focal strain abundance and total biomass with negative binomial errors and a log link were fit in a Bayesian framework using the R package *brms* v2.12.0 [7]. Predictors in both models included community (presence/absence), treatment (antibiotic free, rifampicin and polymyxin) and donor community/slurry (three levels) and the interaction between community presence and donor, and between community presence and treatment. We used default (uninformative) priors, and ran four chain for 2,000 iterations with a warm-up period of 1,000 iterations each, for a combined total of 4,000 samples. We checked model convergence by visualising traceplots and checking diagnostics (Rhat = ∼1).

16S RNA compositional data was analysed using the R packages *Phyloseq* v1.30.0 [8]. We first excluded any OTUs with a total read abundance across all samples of less than five. To visualise compositional similarity across samples (Fig 2b), we performed non-metric multidimensional scaling based on the Bray-Curtis distance between samples.

#### Data availability

16S rRNA Amplicon sequencing data is deposited in the European Nucleotide Archive under the Project accession number PRJEB38496.

#### Supplementary Figure S1-S2

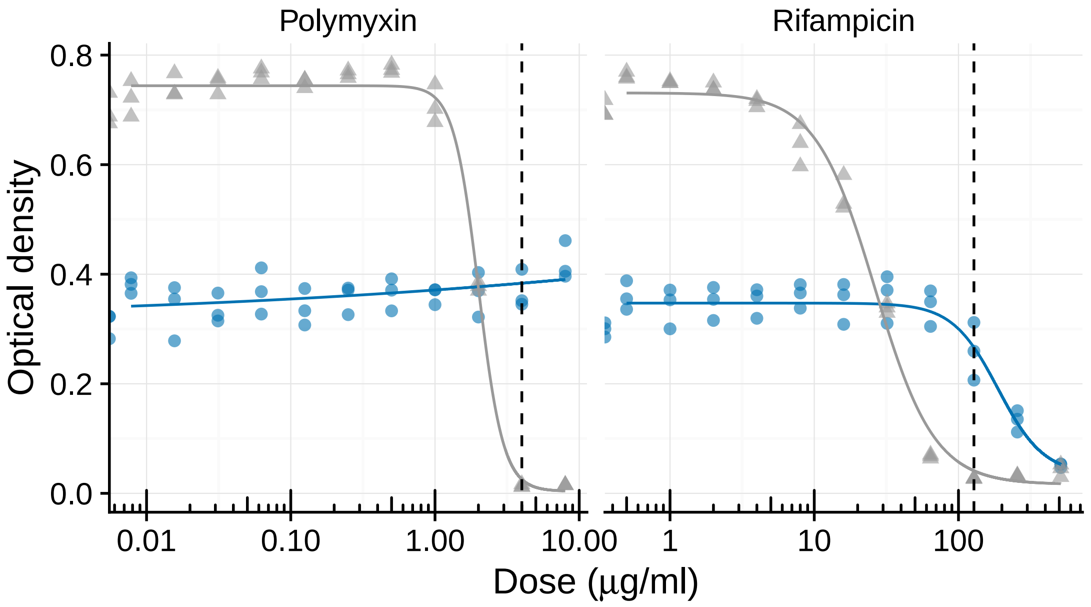


**Figure S1:** Optical density of the focal strain (blue) and its drug sensitive ancestor (grey) after growing for 24 hours in LB supplemented with different concentrations of polymyxin or rifampicin. Vertical dashed lines indicate antibiotic concentrations used in main experiment. Experimental concentrations chosen such that the focal strain can be expected to have an advantage over other sensitive strains in the microbiota.

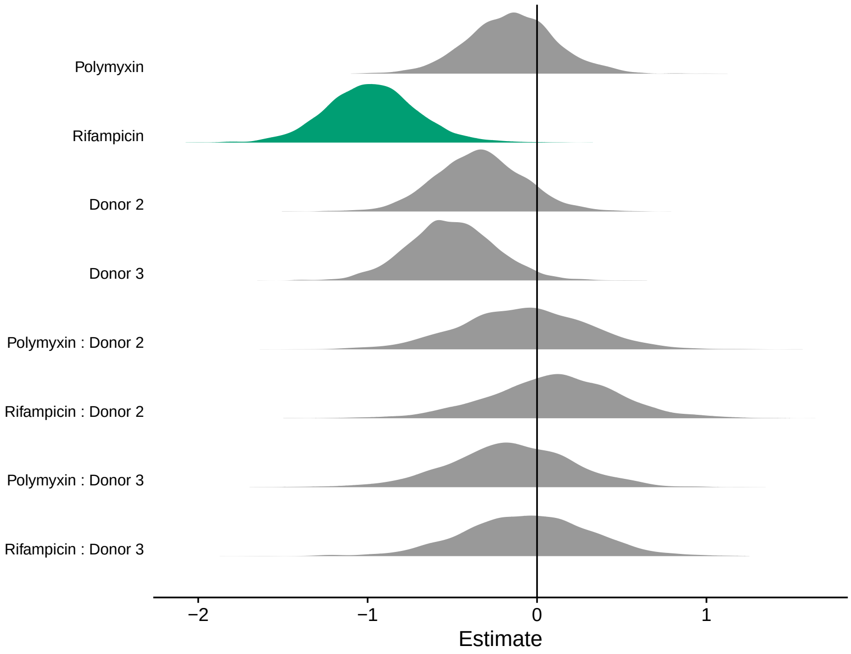


**Figure S2:** Model coefficients (posterior distributions) from a linear model (negative binomial errors) of total bacterial abundance as a function of antibiotic, donor, and their interaction. Posteriors in green have 95% credible intervals that do not overlap with 0 (i.e., there is less than 5% probability there is no effect of the variables/interactions captured by these coefficients). Intercept (not shown) = Donor 1 in the no antibiotic treatment.

### References


[1] Baumgartner, M., Bayer, F., Pfrunder-Cardozo, K. R., Buckling, A. & Hall, A. R. Resident microbial communities inhibit growth and antibiotic-resistance evolution of Escherichia coli in human gut microbiome samples. *PLOS Biology* **18**, e3000465 (2020).

[2] Fouhy, F. *et al.* The Effects of Freezing on Faecal Microbiota as Determined Using MiSeq Sequencing and Culture-Based Investigations. *PLOS ONE* **10**, e0119355 (2015).

[3] Hamilton, M. J., Weingarden, A. R., Sadowsky, M. J. & Khoruts, A. Standardized frozen preparation for transplantation of fecal microbiota for recurrent Clostridium difficile infection. *American Journal of Gastroenterology* **107**, 761–767 (2012).

[4] Robinson, C. D., Auchtung, J. M., Collins, J. & Britton, R. A. Epidemic Clostridium difficile strains demonstrate increased competitive fitness compared to nonepidemic isolates. *Infection and Immunity* **82**, 2815–2825 (2014).

[5] Klümper, U. *et al.* Selection for antimicrobial resistance is reduced when embedded in a natural microbial community. *The ISME Journal* 1–11 (2019).

[6] Klindworth, A. *et al.* Evaluation of general 16S ribosomal RNA gene PCR primers for classical and next-generation sequencing-based diversity studies. *Nucleic Acids Research* **41**, e1 (2013). URL /pmc/articles/PMC3592464/?report=abstracthttps://www.ncbi.nlm.nih.gov/pmc/articles/PMC3592464/.

[7] Bürkner, P.-C. brms: An R package for Bayesian multilevel models using Stan. *Journal of Statistical Software* **80**, 1–28 (2017).

[8] McMurdie, P. J. & Holmes, S. phyloseq: an r package for reproducible interactive analysis and graphics of microbiome census data. *PloS one* **8** (2013).
